# Identification of Iridoid Synthases from *Nepeta* species: Iridoid cyclization does not determine nepetalactone stereochemistry

**DOI:** 10.1101/179572

**Authors:** Nathaniel H. Sherden, Benjamin Lichman, Lorenzo Caputi, Dongyan Zhao, Mohamed O. Kamileen, C. Robin Buell, Sarah E. O’Connor

## Abstract

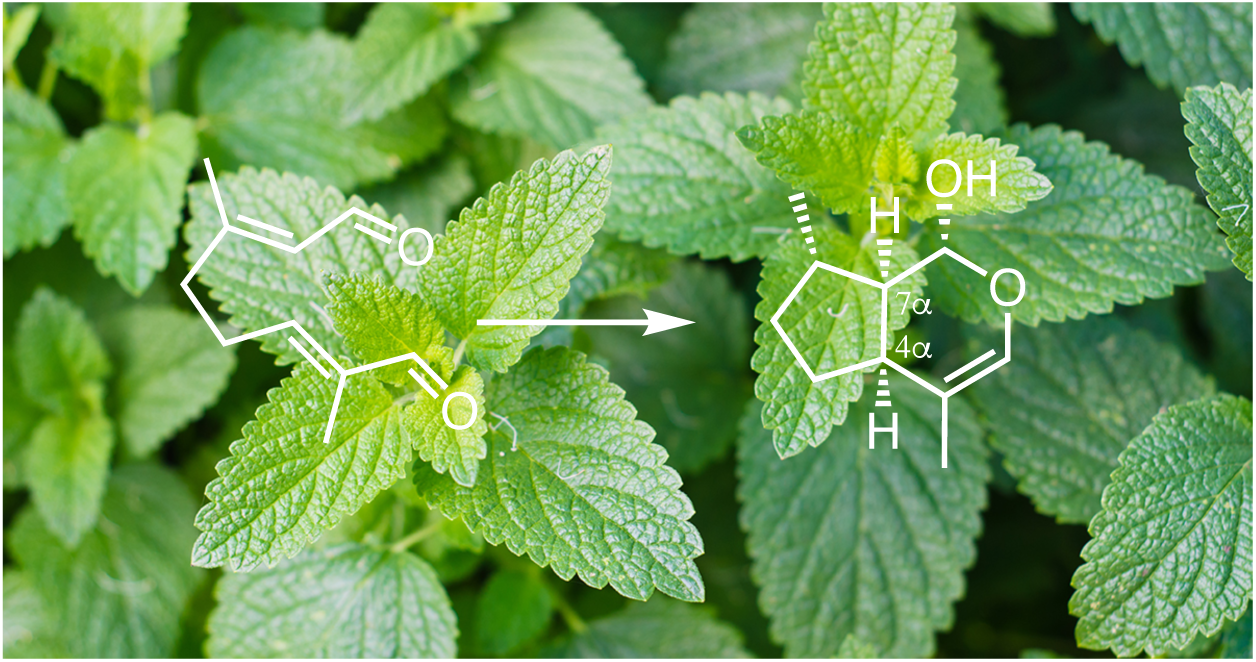
Iridoid synthase from *Nepeta cateria* (catnip) and *Nepeta mussinii*, have been cloned and characterized.

**Abstract:** Nepetalactones are iridoid monoterpenes with a broad range of biological activities produced by plants in the *Nepeta* genus. However, none of the genes for nepetalactone biosynthesis have been discovered. Here we report the transcriptomes of two *Nepeta* species, each with distinctive profiles of nepetalactone stereoisomers. As a starting point for investigation of nepetalactone biosynthesis in *Nepeta*, these transcriptomes were used to identify candidate genes for iridoid synthase homologs, an enzyme that has been shown to form the core iridoid skeleton in several iridoid producing plant species. Iridoid synthase homologs identified from the transcriptomes were cloned, heterologously expressed, and then assayed with the 8-oxogeranial substrate. These experiments revealed that catalytically active iridoid synthase enzymes are present in *Nepeta,* though there are unusual mutations in key active site residues. Nevertheless, these enzymes exhibit similar catalytic activity and product profile compared to previously reported iridoid synthases from other plants. Notably, four nepetalactone stereoisomers with differing stereochemistry at the 4*α* and 7*α* positions – which are generated during the iridoid synthase reaction – are observed at different ratios in various *Nepeta* species. This work strongly suggests that the variable stereochemistry at these 4*α* and 7*α* positions of nepetalactone diastereomers is established further downstream in the iridoid pathway in *Nepeta*. Overall, this work provides a gateway into the biosynthesis of nepetalactones in *Nepeta*.

**Highlights:**

- Species within the *Nepeta* genus (such as catnip) produce nepetalactone iridoids
- The enzymes that produce the iridoid scaffold of nepetalactone were identified from two species of *Nepeta*
- The iridoid synthase enzymes are not responsible for the stereochemical variation in these iridoids

## 1. Introduction

*Nepeta* is a large genus of approximately 250 species within the Lamiaceae (Nepetoideae) family (1,2). *Nepeta* species are well-known for production of nepetalactone (3), the molecule responsible for cat euphoria as exemplified in *Nepeta cataria* (catnip) and other *Nepeta* species (4,5). Notably, nepetalactones are also produced by a number of insects, most notably aphids where they act as sex pheromones (6-13). Not surprisingly, nepetalactone biosynthesis in plants impacts plant-insect interactions, raising the possibility that nepetalactones could be utilized as crop protection agents (14-21).

Early biosynthetic work suggested that nepetalactones are a specific type of monoterpene known as iridoids (22-26). In iridoid biosynthesis in the medicinal plant *Catharanthus roseus*, geranyl-pyrophosphate (GPP) is converted to geraniol by geraniol synthase (GES) (27), hydroxylated to form 8-hydroxygeraniol by geraniol 8-hydroxylase (G8H) (24,28) and then oxidized to 8-oxogeranial by the 8-hydroxygeraniol oxidoreductase (8HGO) (29) (**Figure 1A**). A non-canonical monoterpene synthase, iridoid synthase (ISY) (30), then converts 8-oxogeranial to nepetalactol, along with the open form of this lactol, iridodial. In *Nepeta*, to complete the nepetalactone biosynthetic pathway, an oxidoreductase is believed to convert the nepetalactol to the lactone (31) (**Figure 1B**). The genes that encode GES, G8H, 8HGO, and ISY have been cloned from several plant species, though none have been cloned from a *Nepeta* species, and the gene for the final oxidoreductase to form the lactone has not been identified to date. The stereochemical variation of the nepetalactones is an intriguing aspect of this biosynthetic pathway. Four nepetalactone stereoisomers with differing stereochemistry at the 4*α* and 7*α* positions are observed at different ratios in various *Nepeta* species (referred to as *cis-cis*; *cis-trans*; *trans-cis*; and *trans-trans*; **Figure 1B**) (32-34). The ratio of the stereoisomers appears to play a role in the plants’ ability to deter insects (16,35). However, it is not known how the stereochemistry of these centers is set during the biosynthesis of these compounds. The stereochemistry of the 4*α* and 7*α* carbons are established during the cyclization of 8-oxogeranial to nepetalactol. Iridoid synthase (ISY), the enzyme that catalyzes this cyclization reaction to nepetalactol from the starting substrate 8-oxogeranial, could therefore play a role in this stereochemical variation. Alternatively, downstream isomerase enzymes could be responsible for this stereochemical variation. As a first step to understanding the nepetalactone pathway, we set out to discover and characterize the ISYs from *Nepeta* (Figure 1B).

**FIGURE 1.**
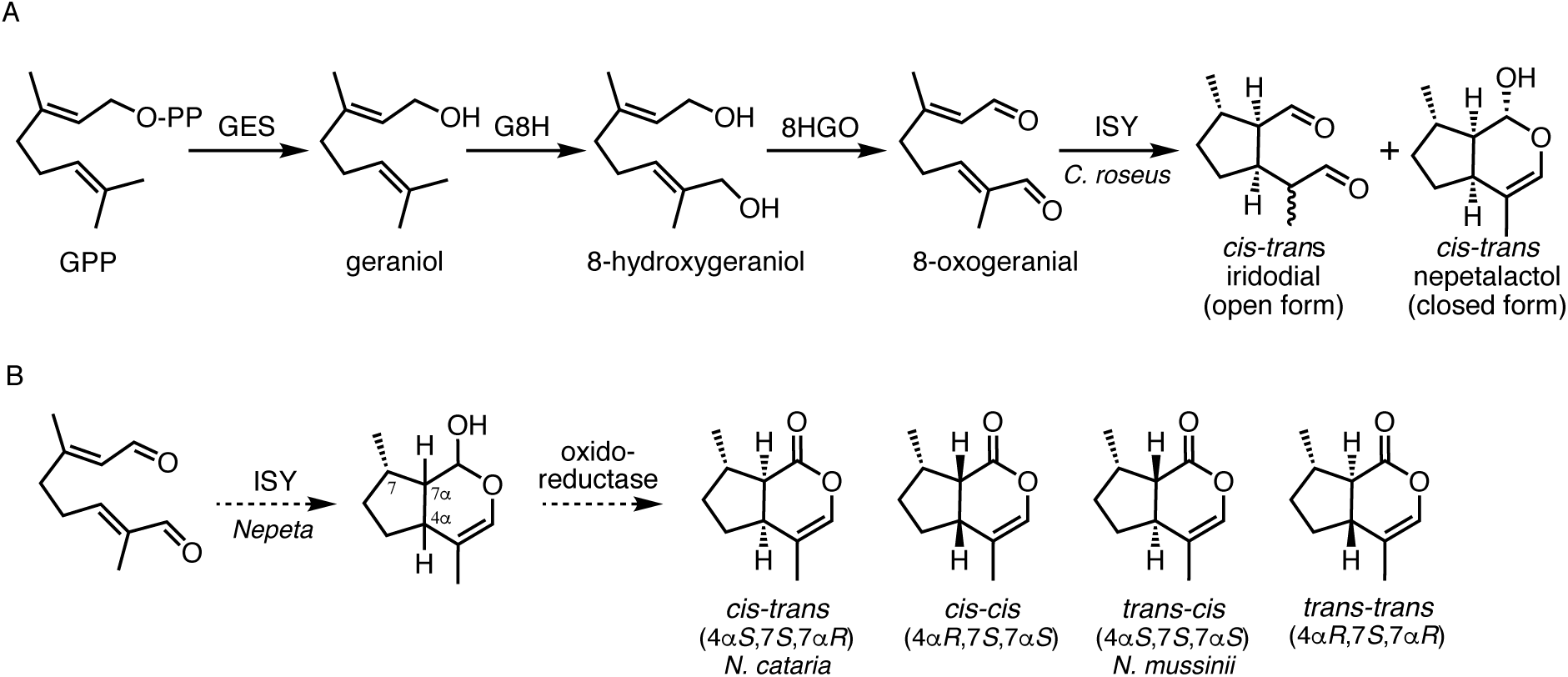
Iridoid biosynthesis. **A.** Formation of the open (iridodial) and closed (nepetalactol) forms of the iridoid skeleton from geranyl pyrophosphate (GPP). GES, geraniol synthase; G8H, geraniol-8-hydroxylase; 8HGO, 8-hydroxygeraniol oxidoreductase; ISY, iridoid synthase. **B.** Nepetalactone biosynthesis entails the conversion of 8-oxogeranial to nepetalactol by ISY as shown in panel A, and an uncharacterized oxidoreductase oxidizes nepetalactol to nepetalactone. Four nepetalactone diastereomers are found in *Nepeta*, with differing stereochemistry at the 4*α* and 7*α* carbons. This study used a cultivar of *N. cataria* that produced primarily *cis-trans-*nepetalactone, and a cultivar of *N. mussinii* that produced primarily *trans-cis*-nepetalactone.

In this study, we report the identification of two *Nepeta* species, *N. cataria* and *N. mussinii*, that produce different profiles of nepetalactone diastereomers in leaf tissue: *N. cataria* produced primarily the *cis-trans* isomer, while *N. mussinii* produced primarily the *trans-cis* isomer (**Figure 1B)**. Leaf tissue from both species was subjected to a transcriptomic analysis, to provide a resource for understanding the molecular basis for the biosynthesis of these compounds. Candidate ISY enzymes were identified by searching for *C. roseus* ISY homologs in the transcriptomes. These genes were then cloned, heterologously expressed and functionally characterized by an *in vitro* assay. Assays of these *Nepeta* ISY (NISY) enzymes with 8-oxogeranial revealed that both *Nepeta* species each harbored two ISY homologs, one of which was substantially more active than the other. These enzymes produced product profiles that were identical to the product profile of the previously reported ISY from *Catharanthus roseus*, an enzyme known to produce the *cis-trans* stereoisomer. These results strongly suggest that the variable stereochemistry of the nepetalactone diastereomers is not established during the cyclization reaction catalyzed by ISY, but that a downstream epimerization step must occur. Notably, the *Nepeta* ISY homologs show surprisingly low sequence identity, as well as changes in the active site residues, to previously characterized ISY enzymes. This discovery highlights the plasticity of genes and encoded enzymes responsible for the synthesis of the iridoid scaffold.

## 2. Results

### 2.1 Selection of plant material

Different species and cultivars of *Nepeta* produce different profiles of nepetalactone diastereomers (16,33). We obtained local *N. cataria* and *N. mussinii* plants and screened the leaves of these plants by GC-MS for production of nepetalactone diastereomers (**Figure 2**). We found that *N. mussinii* plants produced primarily the *trans-cis* (4*αS*,7*S*,7*αS*) lactone, while *N. cataria* produced primarily the *cis-trans* (4*αS*,7*S*,7*αR*) isomer as evidenced by co-elution with authentic standards on a GC-MS (**Figure 2**). Both *N. cataria* and *N. mussinii* have been reported to produce additional nepetalactone isomers, but the nepetalactone profile likely varies widely from cultivar to cultivar. A third *Nepeta* species, *N. racemosa*, was also screened (**Figure 2**). This species showed a product profile identical to *N. cataria* and was not investigated further.

**FIGURE 2.**
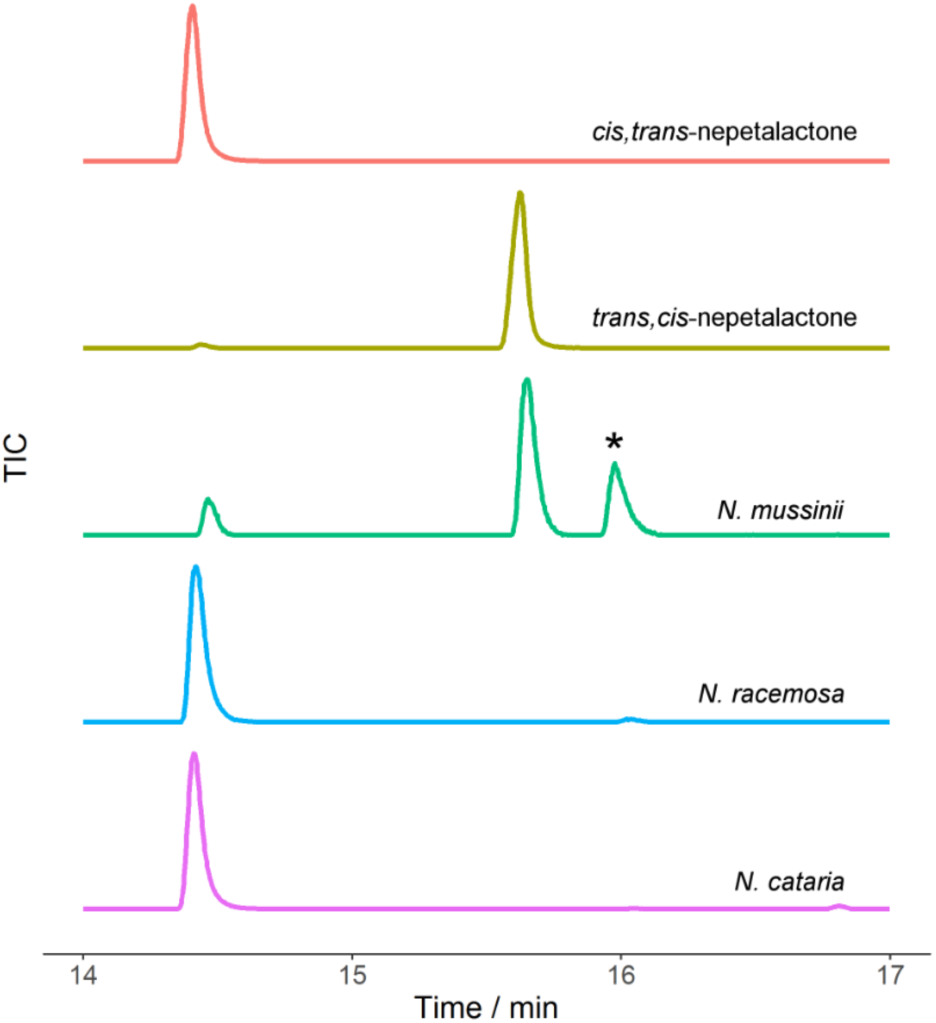
GC-MS chromatograms of leaf tissue extract of three *Nepeta* species. *N. cataria* produced primarily *cis-trans*-nepetalactone, while *N. mussinii* produced primarily *trans-cis*-nepetalactone. A third *Nepeta* species, *N. racemosa*, was also analyzed but was found to have the same chemical composition as *N. cataria*, and was not analyzed further. Authentic standards of *cis-trans* and *trans-cis*-nepetalactones are shown for comparison. One representative GC-MS chromatogram from each species is shown; three biological replicates showed consistent nepetalactone profiles. The peak highlighted with a star is an unknown isomer of nepetalactone.

### 2.2 Cloning, heterologous expression and biochemical assay of ISY from Nepeta

Transcriptomes were obtained for leaf tissue of *N. mussinii* and *N. cataria*. ISY homologs, which are short chain dehydrogenases, are typically annotated as progesterone 5--reductase, the closest known homolog to this enzyme. Two homologs of ISY, from *C. roseus* (30) and *Olea europaea* (olive)(36), that have been functionally characterized were used to search the *Nepeta* transcriptomes for ISY homologs using BLAST. Both *Nepeta* species each contained two distinct homologs of ISY, which were named family 1 and family 2 (amino acid sequence identity between *N. cataria* family 1 (NcISY1) and *N. cataria* family 2 (NcISY2) was 80%; between *N. mussinii* family 1 (NmISY1) and family 2 (NmISY2) was 79%). *Nepeta* homologs exhibited amino acid sequence identities to the ISY from *C. roseus* ranging from 53-58% (**Figure 3**). All of these proteins are members of the short chain dehydrogenase SDR75U family (http://sdr-enzymes.scilifelab.se/).

**FIGURE 3.**
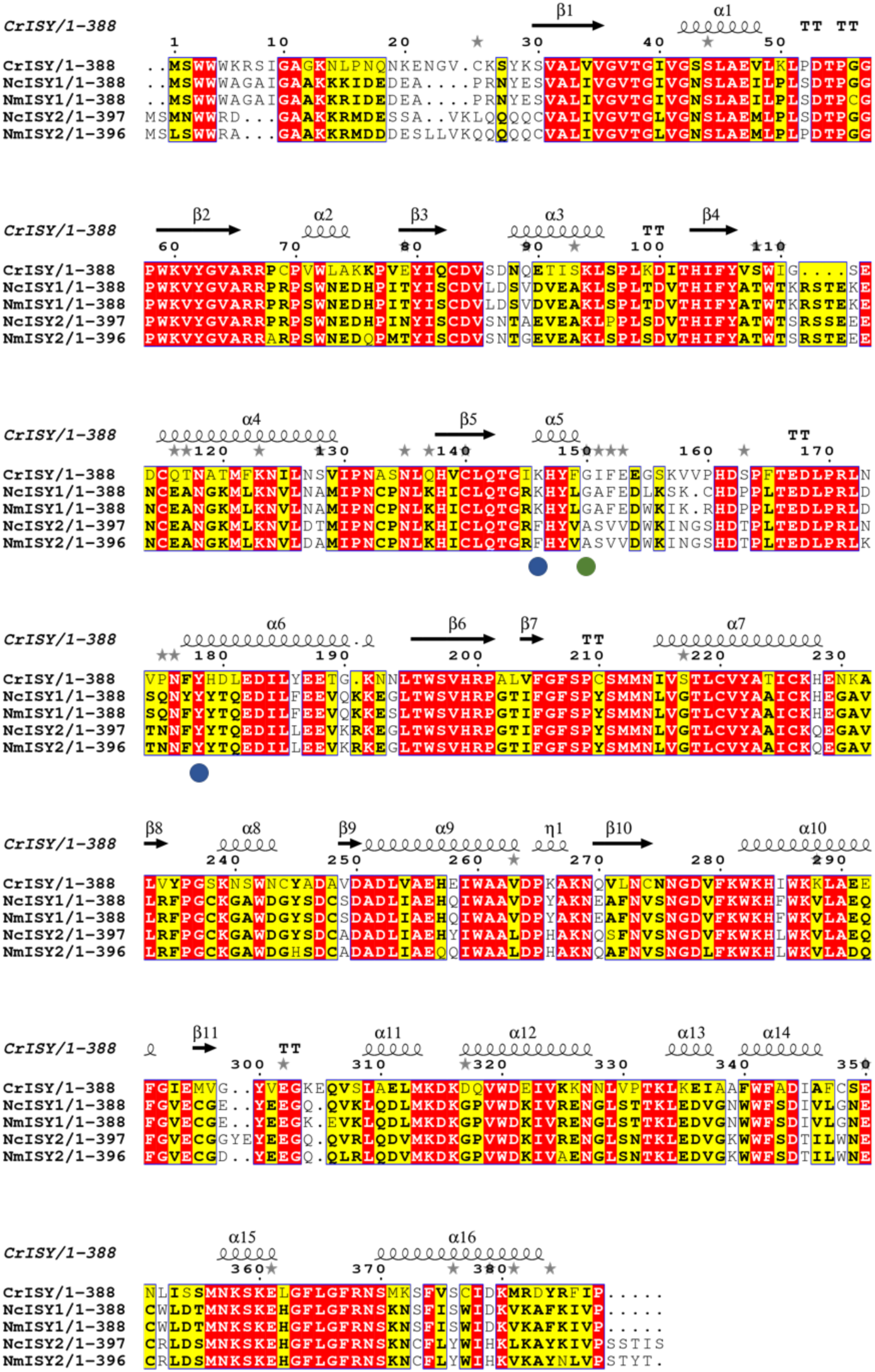
ISY homologs. Alignment of the ISY homologs that were cloned from *Nepeta* compared to ISY from *C. roseus*.

The *Nepeta* ISY enzymes were heterologously expressed in *E. coli* for biochemical characterization, though despite substantial optimization, these enzymes could not be purified to complete homogeneity (**Figure 4**). The enzymes were incubated with the ISY substrate 8-oxogeranial and NADPH according to previously reported ISY assay conditions, and product formation was monitored by GC-MS (30,37). If the stereochemistry of the 4*α* and 7*α* carbons is set by the ISY catalyzed cyclization, then it would be expected that at least one of the *N. mussinii* ISY enzymes would produce large amounts of *trans-cis* product. Authentic standards of *cis-trans-*nepetalactol and iridodials as well as the *trans-cis*-iridodials (the *trans-cis*-nepatalactol is not stable (38) could be resolved on the GC-MS chromatogram (**Figure 5A**).

**FIGURE 4.**
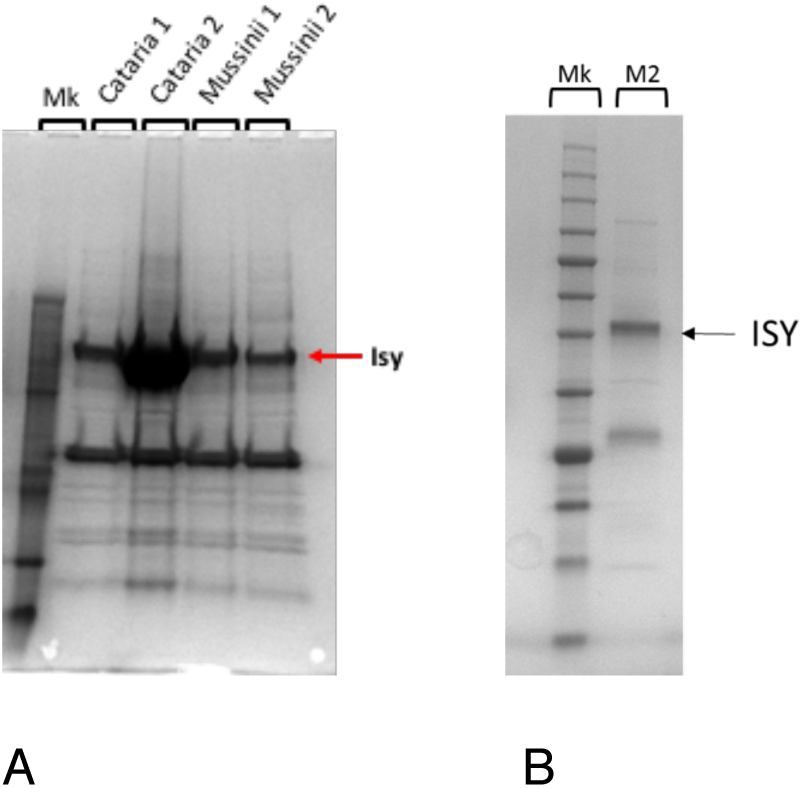
SDS-PAGE of ISY proteins. **A.** All four ISY proteins from *Nepeta* after expression optimization and purification by Ni-NTA chromatography and gel filtration chromatography. Despite extensive optimization of purification conditions (see Experimental), proteins were not entirely homogenous. **B.** SDS-PAGE of *N. mussinii* ISY 2 after Ni-NTA and gel filtration chromatography used for kinetic analysis.

**FIGURE 5.**
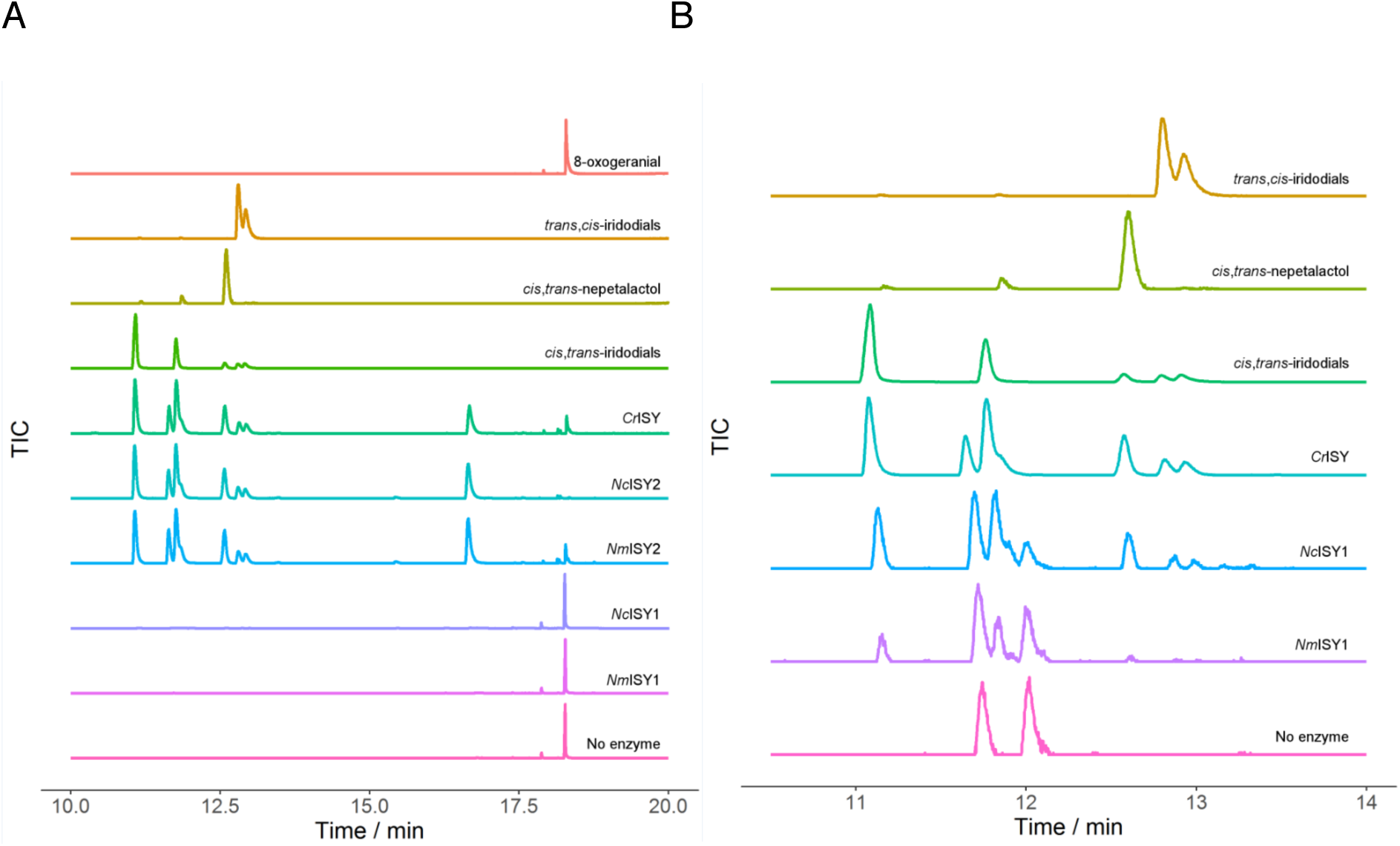
Biochemical assay of the four cloned *Nepeta* ISY enzymes as measured by GC-MS. **A.** GC-MS chromatograms showing authentic standards of substrate 8-oxogeranial, as well as *cis-trans*-nepetalactol, *cis-trans*-iridodial and *trans-cis-*iridodial. Enzymes from *C. roseus*, *N. cataria* (family 2) and *N. mussinii* (family 2) all have nearly identical product profiles. ISY from *N. cataria* (family 1) and *N. mussinii* (family 1) have very low catalytic activity. **B.** Product profiles of ISY from *N. cataria* (family 1) and *N. mussinii* (family 1) compared to that of *C. roseus*. To show trace levels of product, the chromatograms in this panel are normalized to their individual maximum signal intensity within the depicted range.

ISY from *N. cataria* and *N. mussinii* family 1 (NcISY1 and NmISY1) had very low catalytic enzymatic activity as evidenced by an end point assay (**Figure 5A**), and could not subjected to steady state kinetic analysis. The activity was so low that these enzymes are most likely not physiologically relevant in iridoid biosynthesis. However, the trace amount of product that could be detected appeared to be primarily a mixture of *cis-trans-*nepetalactols and iridodials, similar to what is observed with the ISY from *C. roseus* (**Figure 5B**). The family 2 enzymes (NcISY2 and NmISY2) showed substantially higher activity (**Figure 5**). Notably, both NcISY2 and NmISY2 produced a mixture of products identical to the mixture of *cis-trans-*nepetalactol and iridodials produced by the *C. roseus* enzyme.

The steady-state kinetic constants for NmISY2 were obtained, using an assay based on consumption of NADPH (for 8-oxogeranial, K_m_ = 7.3 ± 0.7 μM, k_cat_ = 0.60 ± 0.02 s^-1^; for NADPH, K_m_ = 26.2 ± 5.1, k_cat_ = 0.84 ± 0.06, all data mean±se.) (**Figure 6**). Notably, the availability of the authentic *trans-cis*-iridodial standard demonstrated that small amounts of the *trans-cis* isomer were present in all of the enzymatically catalyzed reactions (**Figure 5A**). However, due to the low levels of the *trans-cis* isomer observed in these assays, it seems unlikely that NcISY2 and NmISY2 provide a direct source of this diastereomer.

**FIGURE 6.**
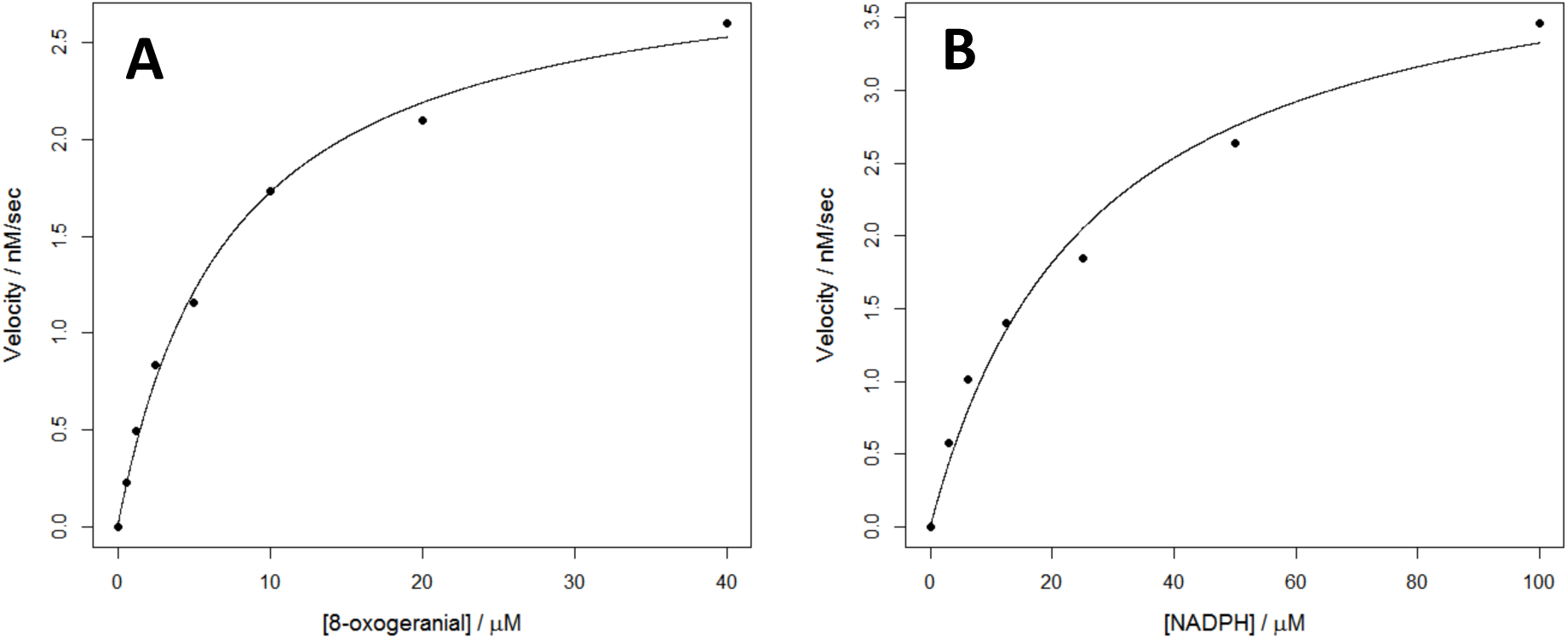
Kinetics of *Nm*ISY2. **A.** Varying 8-oxogeranial concentrations with NADPH concentration at 75 μM. **B.** Varying NADPH concentrations with 8-oxogeranial concentration at 50 μM.

### 2.3 Sequence of Nepeta ISY

Structural and mechanistic studies of ISY from *C. roseus* have revealed that the key active site residue is Tyr178 (37,39). This residue is also conserved in all ISY homologs identified from *Nepeta* (**Figure 3**). A lysine residue in the active site, Lys146, has been shown to play a catalytic role in progesterone 5--reductase, the short chain dehydrogenase most closely related to ISY (40). While this lysine residue is conserved in ISY (*C. roseus*), mutational analysis demonstrated that this residue does not play an essential role in catalysis (37,39). Nevertheless, along with tyrosine, this lysine forms part of the core conserved residues of the short chain dehydrogenase protein family, of which ISY is a member. While family 1 NISY contained this lysine residue, this residue was replaced with a phenylalanine in members of the more catalytically active family 2. Moreover, previous work suggests that Gly150 of ISY (*C. roseus*) allows conformational flexibility of the active site, allowing the enzyme to assume both open and closed forms (37). This residue is mutated to an alanine in family 2 NISY, a mutation that has been shown to lock the active site in an open conformation (37). However, these sequence differences do not appear to alter either the catalytic efficiency or the product profile of these enzymes. This highlights the plasticity of the ISY enzyme responsible for the synthesis of the iridoid scaffold.

## 3. Discussion

Members of the *Nepeta* genus produce nepetalactones, which are iridoid-type monoterpenes that impact plant-insect interactions. The stereochemical variation found among the nepetalactones has a profound influence on the resulting biological activities. However, the biosynthetic pathway of the nepetalactones at the onset of this work was unknown.

Here, we report the transcriptomes of leaves from two *Nepeta* plants, *N. cataria* (catnip) and *N. mussinii* (catmint), plants that produce qualitatively different profiles of nepetalactone stereoisomers. Using this transcriptome database, we searched for homologs of ISY, a key iridoid biosynthetic enzyme that generates the iridoid bicyclic scaffold, along with the 4*α* and 7*α* carbon stereocenters, from the linear substrate 8-oxogeranial. After identification, cloning and heterologous expression of these homologs, *in vitro* biochemical assays indicate that both *Nepeta* species harbor two ISY homologs. Although one homolog (NcISY1 and NmISY1) has only trace catalytic activity, the other (NcISY2 and NmISY2) has robust catalytic activity toward the 8-oxogeranial substrate. Notably, this ISY in both *N. cataria* and *N. mussinii* displayed an unexpected mutation of the active site lysine, as well as mutation of a glycine known to impact the conformational flexibility of the active site.

Surprisingly, the *Nepeta* ISY enzymes assayed gave identical product profiles (**Figure 5**), despite the fact that the stereochemistry of the iridoid scaffolds of these two species are different (**Figure 2**). The product profile of the *Nepeta* enzymes was also very similar to the products produced by the previously reported ISY from *C. roseus*, a plant that produces iridoids exclusively from the *cis-trans* nepetalactol isomer. This discovery highlights that the *Nepeta* ISY enzymes, along with the amino acid sequence differences observed in these enzymes, are not responsible for setting the stereochemistry at the 4*α* and 7*α* carbons. Downstream enzymes that catalyze isomerization of nepetalactol or nepetalactone could be required to yield the stereochemical variation that is observed in the *Nepeta* nepetalactones. The transcriptomic data reported here will facilitate the identification of these downstream enzymes and lead to a greater understanding of the stereochemical variation of these important iridoids produced in *Nepeta*.

## 4. Experimental

### 4.1 Plant material

*Nepeta mussinii* and *N. cataria* were obtained from Herbal Haven, Coldhams Farm, Rickling, Saffron Walden, CB11 3YL, UK. *N. racemosa* ‘Walker’s Low’ was obtained from Notcutts Garden Centre, Daniels Road, Norwich, Norfolk, NR4 6QP, UK. RNA was harvested from mature leaves of plants in a vegetative state using Qiagen RNeasy Plant Mini kit according to the manufacturer’s instructions. For metabolic profiling, approximately 50 mg of frozen leaf tissue were homogenized with a tungsten bead in a ball mill (27 Hz, 30 seconds, 3 repetitions). MeOH (300 μL) was added, mixed vigorously and the resulting slurry was extracted with hexane (600 μL). The top hexane layer was removed, and applied to a Phenomenex Strata SI-1 Silica column (55 μM, 70 Å, 100 mg/mL). The hexane flow through was discarded. The compounds of interest were eluted from the column with 20% EtOAc/Hexane (500 μL), and this eluant was analysed by GC-MS. Three individual plants were analysed for each species; no significant metabolite variation between individuals was observed.

### 4.2 Compounds

Syntheses and isolation of key compounds are outlined below. All of the compounds have been described previously. ^^1^^H NMR spectra were used to demonstrate compound purity (Supporting Information). 8-oxogeranial was synthesized from geranyl acetate as previously described in Geu-Flores *et al*. *Cis*-*trans*-nepetalactol was synthesized by DIBAL-H reduction of *cis*-*trans*-nepetalactone as previously described in Geu-Flores *et al*.

#### Cis-trans-nepetalactone

Leaves and shoots (∼ 100 g) of *Nepeta fassinii* “Six Hills Giant” (Burncoose Nurseries, Gwennap, Redruth, Cornwall, TR16 6BJ, UK) and water (300 mL) were blended (3 x 1 min) in a kitchen blender. The dark green slurry was extracted with dichloromethane (10 x 100 mL). Fractions were analysed by GC-MS to determine nepetalactone presence. Combined fractions were filtered, washed with brine (200 mL), dried over anhydrous Na_2_SO_4_ and evaporated to dryness *in vacuo* yielding a yellow oil (1.098 g crude). The nepetalactone was purified by silica flash chromatography (60 g silica, 3x25 cm). The crude oil was loaded onto the column in hexane and eluted in hexane/EtOAc fractions (40 mL) with increasing concentrations of EtOAc. Fractions were analysed by GC-MS; those containing the desired nepetalactone isomer were combined and evaporated to dryness yielding the pure *cis*-*trans*-nepetalactone (155 mg). Product identity was verified by NMR spectroscopy.

#### Trans-cis-nepetalactone

Catnip oil (2 mL, Health and Herbs, 425 Ellsworth St. SW Albany, OR 97321, USA, https://betterhealthherbs.com/) was purified by silica flash chromatography. The crude oil was applied to the column and eluted with hexane/EtOAc with increasing concentrations of EtOAc. Fractions containing the same compounds (as analysed by TLC) were combined and evaporated to dryness. Pure β-caryophyllene (700 mg) and *trans*-*cis*-nepetalactone (441 mg) were isolated. Compound identity was determined by NMR spectroscopy and comparison to literature values.

#### Trans-cis-iridodials

*Trans-cis*-nepetalactone (212 mg) was dissolved in hexanes (30 mL) and cooled to -78 °C in an inert atmosphere. A solution of DIBAL-H in hexanes (213 mg, 5% v/v) was added dropwise over 10 minutes and left to stir for 1 hour. Baeckstrom reagent (1.6 g, celite:Na_2_SO_4_ 1:1) was added and the reaction was stirred for 1 hour at -78 °C and then 2 hours at room temperature. The mixture was filtered through a glass frit and concentrated *in vacuo* yielding the crude product. The compound was purified by silica flash chromatography and eluted in EtOAc/hexane with increasing concentrations of EtOAc. Fractions were combined and evaporated to dryness to yield a mixture of the two *trans*-*cis*-iridodials (64 mg). The identity of the compound was verified by 1H NMR.

#### Cis-trans-iridodials

*Cis-trans*-nepetalactol (0.6 mM) was incubated in dilute aqueous HCl (100 mM) overnight, extracted into EtOAc and analysed by GC-MS.

### 4.3 Transcriptome and bioinformatics

RNA-sequencing (RNA-seq) libraries were constructed using the Kappa stranded RNA-seq kit and libraries were sequenced on an Illumina HiSeq 2500 generating 150 nt paired-end reads. Read quality was assessed using FASTQC (v0.11.2; http://www.bioinformatics.babraham.ac.uk/projects/fastqc/) with default parameters and adaptors and low quality sequences removed using Trimmomatic (Parameter: LEADING:10 TRAILING:10 SLIDINGWINDOW:4:15 MINLEN:30) (v0.32, (41). Only surviving paired-end reads were used to generate *de novo* assemblies using Trinity (v2014-07-17) (42) and only transcripts >250 bp were reported (**Table 1**). The protein sequences of iridoid synthases from *Catharanthus roseus* (AFW98981.1) and *Olea europaea* L (ALV83438.1) were used to BLAST the predicted peptides of the *N. mussinii* and *N. cataria* transcriptomes (43). Two ISY candidates from each transcriptome were identified.

**Table 1.**
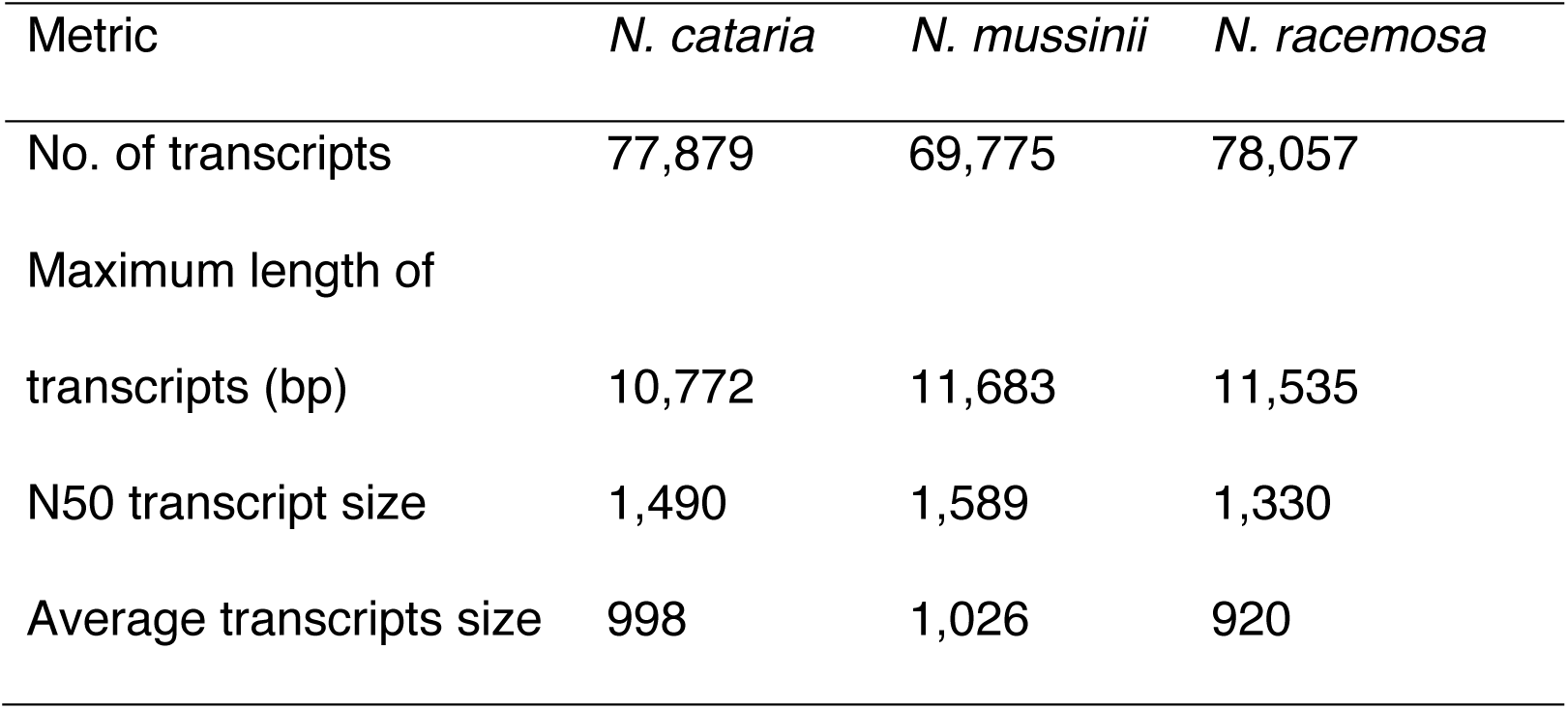
Transcriptomic metrics.

| Metric | *N. cataria* | *N. mussinii* | *N. racemosa* |
| --- | --- | --- | --- |
| No. of transcripts | 77,879 | 69,775 | 78,057 |
| Maximum length of transcripts (bp) | 10,772 | 11,683 | 11,535 |
| N50 transcript size | 1,490 | 1,589 | 1,330 |
| Average transcripts size | 998 | 1,026 | 920 |

### 4.4 Cloning and protein expression

cDNA was prepared from leaf RNA using Invitrogen SuperScript III First-Strand Synthesis System kit following the manufacturer’s instructions. Primers were designed based on the ISY transcript sequences and with 5’-overhangs for cloning (**Table 2**). These were used to PCR amplify ISY genes from the cDNA and clone directly into a pOPINF expression vector using an InFusion HD cloning kit. The pOPINF vector encodes an N-terminal His-tag. The gene sequences were verified by Sanger sequencing. Two ISY candidates from *N. mussinii* were obtained (*Nm*ISY1 and *Nm*ISY2). Sequencing revealed that three ISY candidates were cloned from *N. cataria*. Two were cloned from with same primer pair (*Nc*ISY1) and had 97% amino acid identity, and thus, only one was investigated further. The two ISY candidates from *N. cataria* investigated further were named *Nc*ISY1 and *Nc*ISY2. Proteins were subjected to a variety of expression conditions to maximize expression, with the most optimal conditions described below. All genes were expressed in strain soluBL21 (DE3) (Genlantis). Cells were pre-cultured overnight at 37 °C in LB medium containing 100 μg/mL carbenicillin. Aliquots of 200 μL each were used to inoculate two 2-L flasks containing 2YT medium and carbenicillin. When cultures reached an OD600 of 0.5–0.8 after ∼6 hours shaking at 37 °C, protein production was induced by adding IPTG at a final concentration of 500 μM. Protein was expressed at 18 °C for ∼16 h. Cells were harvested by centrifugation, the supernatant was discarded and pellets were resuspended in 100 mL of 50 mM Tris-HCl buffer (pH 7.0) containing 300 mM NaCl, 1 mM DTT, one tablet of Complete EDTA free protease inhibitor (Roche), and 0.2 mg/mL lysozyme. The cells were disrupted by 7 minutes sonication on ice in cycles of 2 seconds sonication followed by a 3 second break. All subsequent steps were conducted at 4 °C. The lysate was centrifuged for 20 minutes at 35,000 *g* and the supernatant containing the soluble protein was applied on a 5 mL HisTrap column connected to an Äkta Xpress purifier (GE healthcare). His-tagged protein was eluted with a step gradient of 20 mM to 500 mM imidazole in 50 mM Tris/glycine buffer adjusted to pH 8.0 with hydrochloric acid and supplemented with 5% (v/v) glycerol, 0.5 M NaCl, and 1 mM DTT. Fractions containing the protein of interest were collected and concentrated to 2–3 mL in an Amicon 10 kDa MWCO centrifugal filter (Millipore). The concentrated solution was further purified by size-exclusion chromatography on a Superdex 200 16/60 GF column (GE Healthcare). Fractions corresponding to the molecular weight of the ISY (*C. roseus*) dimer (83 kDa) were collected, combined, concentrated in an Amicon 10 kDa MWCO centrifugal filter and stored at -20 °C after flash freezing in liquid nitrogen. Protein concentrations were determined spectrophotometrically at 280 nm using extinction coefficients calculated with the protparam tool (http://web.expasy.org/protparam/). Despite extensive optimization of purification conditions, it was not possible to purify proteins to complete homogeneity as judged by SDS-PAGE (Figure 4).

**Table 2.**
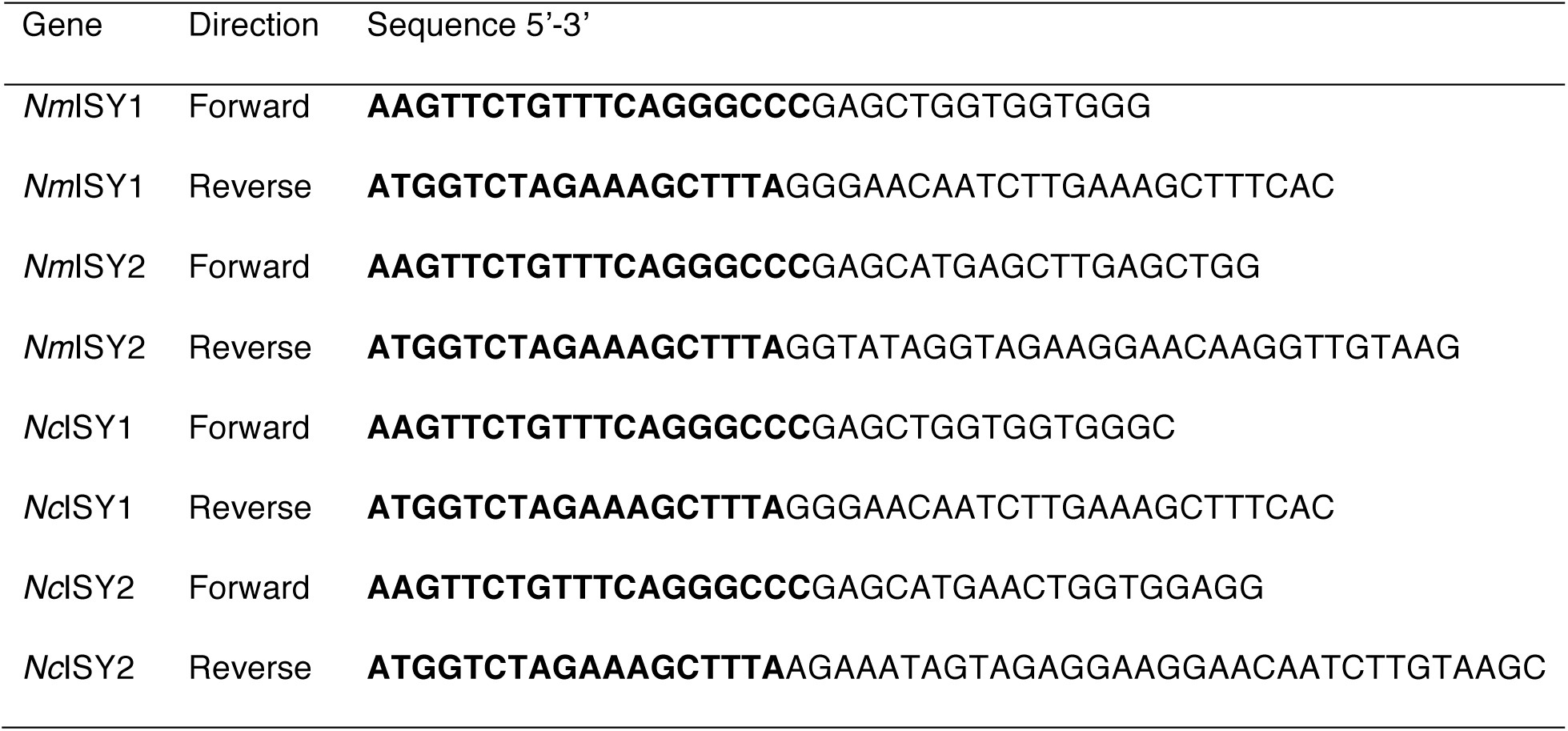
Primers used for cloning. Overhangs used for pOPINF InFusion are in bold.

### 4.5 Assay conditions, end point reactions

Reactions were conducted with 8-oxogeranial (500 μM), NADPH (1 mM), MOPS pH 7.0 (200 mM), NaCl (100 mM), THF (0.5% v/v) and *Nepeta* ISYs (0.5 μM). A negative control was conducted without enzyme, and a positive control was conducted with ISY (*C. roseus*). Reactions were incubated for 3 hours at 30 °C before extraction with EtOAc (100 μL) and analysis on GC-MS.

### 4.6 Kinetic measurement conditions

Kinetics of NADPH consumption were determined spectrophotometrically on a PerkinElmer Lambda 35 instrument at a wavelength of 340 nm in cuvettes with 1 cm path length. Reactions contained 200 mM MOPS buffer pH 7.0, 100 mM sodium chloride and 5 nM *Nm*ISY2, in a total volume of 800 μL. 8-oxogeranial was added from a stock solution in inhibitor free THF. A THF co-solvent concentration of 1-1.4% was maintained in the assay to ensure substrate solubility. For determination of 8-oxogeranial parameters, 75 μM NADPH was used. For determination of NADPH kinetic parameters, 50 μM 8-oxogeranial was used. Cuvettes were equilibrated to 25 °C before the reaction was initiated by addition of enzyme. Absorbance values were recorded at a rate of 1 Hz. The R software environment was used to fit linear initial rates over 2-5 minutes of the enzyme reaction. Background NADPH consumption was subtracted from initial rates. The Michaelis-Menten equation was fit to data points in R by the nls function to obtain kinetic data. The enzyme *Nm*ISY2 appeared to lose activity during storage at -20 °C.

### 4.7 GC-MS method

Samples were injected in split mode (2 μL, split ratio 20:1) at an inlet temperature of 220 °C on a Hewlett Packard 6890 GC-MS equipped with a 5973 mass selective detector (MSD), and an Agilent 7683B series injector and autosampler. Separation was performed on a Zebron ZB5-HT-INFERNO column (5% phenyl methyl siloxane; length: 35 m; diameter: 250 μm) with guard column. Helium was used as mobile phase at a constant flow rate of 1.2 mL/minute and average velocity 37 cm/s. After 5 minutes at 80 °C, the column temperature was increased to 110 °C at a rate of 2.5 K/min, then to 280 °C at 120 K/min, and kept at 280 °C for another 4 minutes. A solvent delay of 5 minutes was allowed before collecting MS spectra at a fragmentation energy of 70 eV.

## Acknowledgments

We gratefully acknowledge funds from the US National Science Foundation Plant Genome Research Program to CRB and SOC (IOS-1444499) as well the UK Biotechnological and Biological Sciences Research Council (BBSRC) and Engineering and Physical Sciences Research Council (EPSRC) joint-funded OpenPlant Synthetic Biology Research Centre (BB/L014130/1). The raw RNA-sequencing reads used in the transcriptome assembly have been deposited in the National Center for Biotechnology Information Sequence Read Archive under BioProject ID (PRJNA379302). The assembled transcriptomes are available at the Dryad Digital Repository under doi (to be released upon publication). Sequences for NmISY1 (KY882235), NmISY2 (KY882236), NcISY1 (KY882233), NcISY2 (KY882234) have been deposited in GenBank.

## Author contributions

N.H.S. and S.E.O. conceived the study; N.H.S. identified plants, cloned ISY genes, prepared standards, obtained initial results, and generated conclusions; L.C. purified the protein; B.L. generated kinetic data; M.O.K. screened plants; C.R.B and D.Z. generated transcriptome; S.E.O. wrote the manuscript.

